# T-bet^+^ CD11c^+^ B cells are critical for anti-chromatin IgG production in the development of lupus

**DOI:** 10.1101/116145

**Authors:** Ya Liu, Shiyu Zhou, Jie Qian, Yan Wang, Xiang Yu, Dai Dai, Min Dai, Lingling Wu, Zhuojun Liao, Zhixin Xue, Jiehua Wang, Guojun Hou, Jianyang Ma, John B. Harley, Yuanjia Tang, Nan Shen

**Affiliations:** Shanghai Institute of Rheumatology, Renji Hospital, School of Medicine, Shanghai Jiao Tong University, Shanghai, China; Institute of Health Sciences, Shanghai Institutes for Biological Sciences (SIBS) & Shanghai Jiao Tong University School of Medicine (SJTUSM), Chinese Academy of Sciences (CAS), Shanghai, China; Cincinnati Children’s Hospital Medical Center, University of Cincinnati College of Medicine, and Cincinnati VA Medical Center, Cincinnati; Shanghai Institute of Rheumatology, Renji Hospital, School of Medicine, Shanghai Jiao Tong University, Shanghai, China; Institute of Health Sciences, Shanghai Institutes for Biological Sciences (SIBS) & Shanghai Jiao Tong University School of Medicine (SJTUSM), Chinese Academy of Sciences (CAS), Shanghai, China; Center for Autoimmune Genomics and Etiology (CAGE), Cincinnati Children’s Hospital Medical Center, Cincinnati, Ohio, USA; Collaborative Innovation Center for Translational Medicine at Shanghai Jiao Tong University School of Medicine; State Key Laboratory of Oncogenes and Related Genes, Shanghai Cancer Institute, Renji Hospital, Shanghai Jiao Tong University School of Medicine, Shanghai

**Keywords:** Systemic lupus erythematosus, chronic graft-versus-host disease, T-bet; anti-chromatin antibody, B cell

## Abstract

A hallmark of systemic lupus erythematosus is high titers of circulating autoantibody. A novel CD11c^+^ B cell subset has been identified that is critical for the development of autoimmunity. However, the role of CD11c^+^ B cells in the development of lupus is unclear. Chronic graft-versus-host disease (cGVHD) is a lupus-like syndrome with great autoantibody production. In the present study we investigated the role of CD11c^+^ B cells in the pathogenesis of lupus in the cGVHD model. Here, we found the percentage and absolute number of CD11c^+^ B cells and titer of sera anti-chromatin IgG and IgG2a antibody were increased in cGVHD mice. CD11c^+^ plasma cells from cGVHD mice produced large amounts of anti-chromatin IgG2a upon stimulation. Depletion of CD11c^+^ B cells reduced anti-chromatin IgG and IgG2a production. T-bet expression was further shown to be upregulated in CD11c^+^ B cells. Knockout of T-bet in B cells alleviated cGVHD. The percentage of T-bet^+^ CD11c^+^ B cells was elevated in lupus patients and positively correlated with serum anti-chromatin levels. Our findings suggest T-bet^+^ CD11c^+^ B cells contribute to the pathogenesis of lupus and provides potential target for therapeutic intervention.

## Introduction

Systemic lupus erythematosus (SLE) is a prototypic autoimmune disease characterized by an array of autoantibody that targets multiple normal cellular components [1–3]. When encountered with self-antigens, autoantibody can bind with them to form immunoglobulin complexes (ICs), which deposit in the kidney, activate complement system, and appear a series of inflammatory responses. Accumulating evidence has indicated that the presence of certain autoantibody is highly associated with some symptoms of the disease [4, 5]. For example, antiphospholipid antibodies are critically linked to the development of thrombotic events and obstetric morbidity [6], anti-RNP antibodies are associated with myositis and Raynaud’s phenomenon [7], and anti-dsDNA antibodies are associated with lupus nephritis [8]. However, the characterization of B cell that producing the distinct kinds of pathogenic antibodies is still unclear. Chromatin, the native complex of histones and DNA found in the cell nucleus of eukaryotes, consists of approximately 40% DNA, 40% histones, and 20% non-histone proteins, RNA and other macromolecules [9]. Data accumulated in recent years, have indicated that anti-chromatin autoantibody were involved in the pathogenesis of SLE and associated with disease activity and lupus nephritis [10–12]. For example, immunization with active chromatin induced lupus-like syndrome in BALB/c mice [13]. Several studies of lupus-like murine models have found genetic loci, such as sle1, that are related to anti-chromatin antibodies production [14]. In certain knock-out mice (for C1q, serumamyloid P [SAP] and Dnase I), impaired phagocytosis of apoptotic material may result in an autoimmune response to chromatin and glomerulonephritis [15, 16]. However, it is not quite clear how B cells lose itself tolerance to chromatin in the development of lupus.

Transfer of MHC II-mismatched splenocytes from Bm12 mice into B6 mice causes a chronic graft versus host disease (cGVHD), which is characterized by the production of high titers of autoantibody and immunopathology that closely resemble SLE [17–19]. The majority of autoantibody in cGVHD are anti-chromatin and anti-erythrocytes [20]. Recently, a novel B cell subset- CD19^+^ CD11c^+^ cell is identified in aged female B6 mice, which is able to produce large amounts of anti-chromatin autoantibody in response to TLR7 ligand in vitro [21]. However, little is known about the differentiation and function of CD 19^+^ CD11c^+^ cells in the development of lupus.

In this study, we present data identifying T-bet as an essential factor in regulating the development of CD11c^+^ B cells in cGVHD induced lupus. Deletion of T-bet significantly decreased the frequency as well as the absolute number of CD19^+^ CD11c^+^ cells during cGVHD, with reduced circulating levels of anti-chromatin autoantibody. Importantly, T-bet^+^ CD11c^+^ CD19^+^ B cells were remarkably increased in lupus patients and positively correlated with titer of anti-chromatin antibodies. Therefore, taken together, our data demonstrated that T-bet^+^ CD11c^+^ CD19^+^ B cells are critical for the anti-chromatin autoantibody production, which might be explored as a therapeutic target for rectifying the abnormally produced anti-chromatin in SLE

## Patients and methods

### Human study subject samples

22 patients with SLE and 10 Healthy donors were recruited for analyzing the percentage of T-bet^+^ CD11c^+^ CD19^+^ cells in PBMC. All SLE patients were recruited from Renji Hospital and fulfilled the American College of Rheumatology 1982 revised criteria for SLE [22] and 14 of these patients met the ACR criteria for lupus nephritis [23]. The Systemic Lupus Erythematosus Disease Activity Index (SLEDAI) score was determined for each patient at the time of the blood draw [24]. Patients were categorized as having active disease (scores >4) or inactive disease (scores ≤ 4) based on the SLEDAI results. Additional clinical information about the subjects is listed in Table 1. Informed consent was obtained from all of the subjects. The study was approved by the Research Ethics Board of Shanghai Renji Hospital.

**Table 1.**
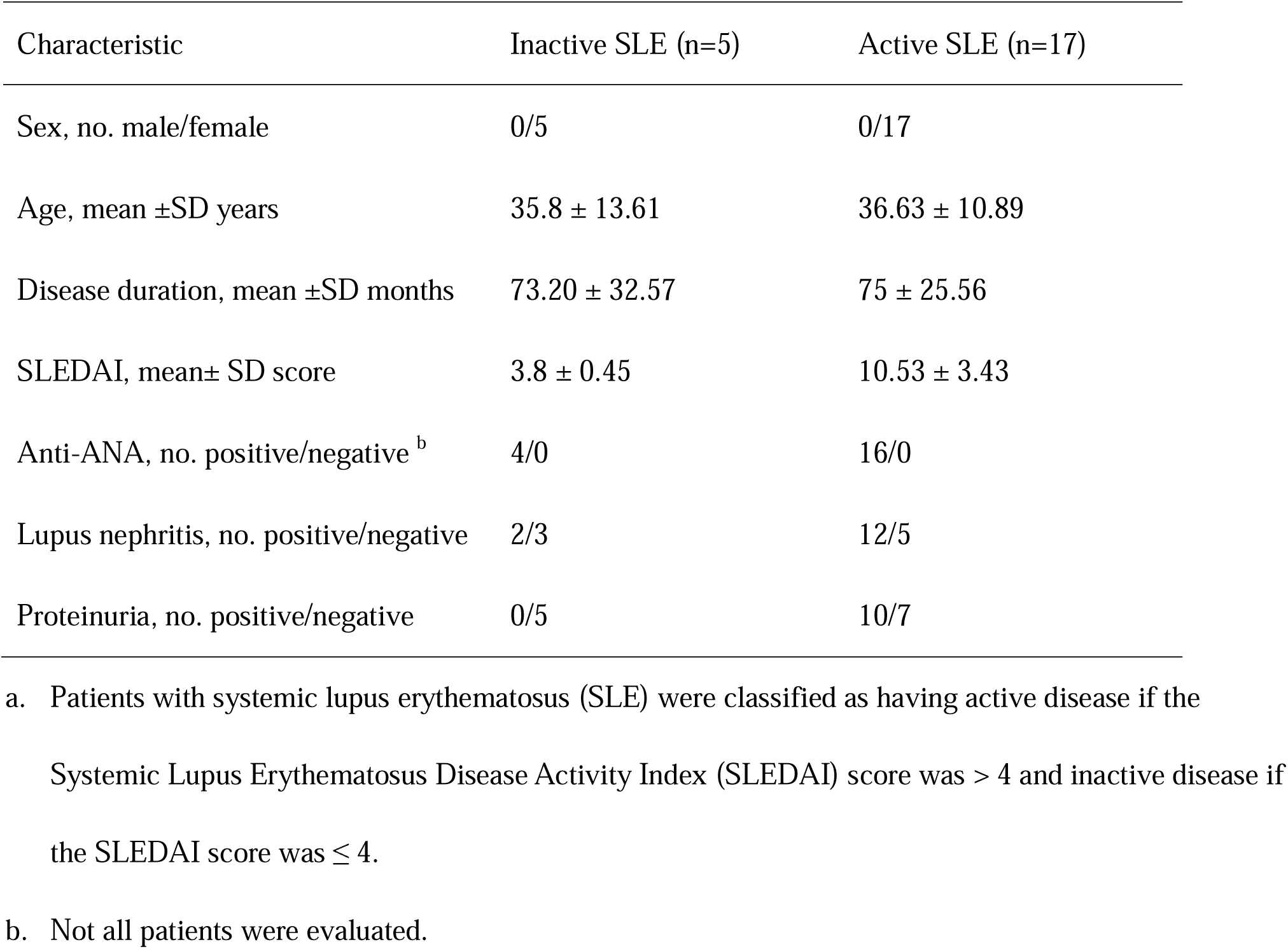
Clinical features of the 18 patients with SLE, by disease activity group^a^.

| Characteristic | Inactive SLE (n=5) | Active SLE (n=17) |
| --- | --- | --- |
| Sex, no. male/female | 0/5 | 0/17 |
| Age, mean ±SD years | 35.8 ± 13.61 | 36.63 ± 10.89 |
| Disease duration, mean ±SD months | 73.20 ± 32.57 | 75 ± 25.56 |
| SLEDAI, mean± SD score | 3.8 ± 0.45 | 10.53 ± 3.43 |
| Anti-ANA, no. positive/negative <sup>b</sup> | 4/0 | 16/0 |
| Lupus nephritis, no. positive/negative | 2/3 | 12/5 |
| Proteinuria, no. positive/negative | 0/5 | 10/7 |

a. Patients with systemic lupus erythematosus (SLE) were classified as having active disease if the Systemic Lupus Erythematosus Disease Activity Index (SLEDAI) score was $> 4$ and inactive disease if the SLEDAI score was $\leq 4$ .
b. Not all patients were evaluated.

### Mice

B6(C)-H2-Ab 1 bm12/KhEgJ (Bm12), C57BL/6J (B6), B6.129S7-Ifngr1tm1Agt/J (B6.IFNGR1-/-), B6.129P2-Igh-Jtm1Cgn/J (μMT), and B6.FVB-Tg (Itgax-DTR/EGFP) 57Lan/J (B6.CD11c-DTR) were purchased from The Jackson Laboratory (Bar Harbor, ME). Bm12 and B6 were propagated in the animal facility at Cincinnati Children’s Hospital Medical Center (CCHMC) (Cincinnati, U.S.A). μMT and B6.CD11c-DTR mice were maintained in the animal facility at the Institute of Health Science (IHS) (Shanghai, China). All animals were 10-to 12-week-old at the time of experimentation. All animal protocols were approved by the Animal Care and Use Committee of CCHMC and IHS.

### cGVHD induction

Single-cell suspension of Bm12 splenocytes was prepared in 1 × PBS, filtered through 0.2 μm sterile nylon mesh. 5 × 10^^^7 splenocytes was then i.p. injected into B6 mice. After two weeks, the recipient mice were sacrificed for analysis.

### Abs and Flow cytometry

The following mAbs used for staining were purchased from BioLegend: Bv421 anti-CD4, PerCP anti-CD8a, APC anti-CD69, PE-Cy7 anti-CXCR5, PE anti-PD-1, AF-700 anti-CD44, FITC anti-CD62L, APC-Cy7 anti-CD 19, PerCP-Cy5.5 anti-CD19, PE anti-FasR, FITC anti-GL7, BV421 anti-CD138, PE anti-CD138, PE-Cy7 anti-CD 11b, APC-eF780 anti-CD11c, APC anti-IFNγ, and APC anti-T-bet.eF506 Live/Dead dye was obtained from eBioscience. Human Antibodies were bought from BD bioscience: Bv605 anti-CD 19, FITC anti-CD 11c. Cells were fixed in BD Cytofix™ buffer (BD bioscience) before FACS analysis. Intracellular staining for T-bet was performed by using BD Cytofix/Cytoperm™ Kit (BD bioscience). Data were collected on Fortessa2 and LSR-II flow cytometer and analyzed by FlowJo software.

### Cell isolation and in vitro culture

Spleen cells from Bm12→B6 cGVHD mice were pooled together (n=5). CD19^+^ B cells were first positively selected by using CD19 positive MACS beads. CD11c^+^ CD138^+^ and CD11c^−^ CD138^+^ cells were then sorted by FACSDiva. Equal cells were then cultured in RPMI-1640 supplemented with 10% FBS, 2mM L-glutamine, 100 U/ml penicillin, 100 U/ml streptomycin, and 50 μM 2-mercaptoethanol for 72 h. 1 ug/mL R848 was used for cell stimulation. Supernatant was then assayed by anti-chromatin Abs ELISA. To test IFNγ + T cells, 2 × 107 spleen cells from Bm12 to B6 cGVHD mice were cultured in the cell culture medium above with 1 × Cell Stimulation Cocktail (plus protein transport inhibitors) (eBioscience) for 6 h. Then stained with Live/Dead dye, anti-CD4, and anti-IFN-γ mAbs.

### ELISA for anti-chromatin Abs

Chromatin was prepared as previously described [25, 26]. Plates were coated with chromatin at 3 ug/ml overnight at 4 °C. The plates were then washed and incubated with blocking buffer for 2 h at Room Temperature (RT). The plates were washed again, and then 1/500 diluted Serum samples were added in duplicate and incubated for 2h at RT. Biotin-labeled anti-IgG, anti-IgG1, anti-IgG2a, anti-IgG2b, and IgG3 (BioLegend) were used as capture antibodies. And Streptavidin-conjugated HRP (Thermo Scientific) was used as detective antibody. 1 × TMB (eBioscience) was then added and incubated for 30 min at RT. 2 N sulfuric acid was used as stop solution. O.D values were then measured at 450 nm and 570 nm.

### Depletion of CD11c^+^ B cell in cGVHD mice

CD11c-DTR mice with cGVHD were induced by transferring 5 × 10^7^splenocytes from Bm12. For depletion of CD11c^+^ cells, mice were then intraperitoneally injected with 100 ng Diphtheria toxin in 200 uL PBS (Sigma) at day 7, day 9 and day 11-post cGVHD induction. The efficiency of depletion was then examined by flow cytometry14 day post cGVHD induction.

### Statistical analysis

Data were analyzed using GraphPad Prism (version 5.01). Statistical differences were calculated using one-way ANOVA and the Student t test. Nonparametric correlation (spearman) was used for correlation studies. Values are presented as the mean ± SD. A value of p<0.05 was considered to be statistically significant.

## Results

### CD11c^+^ B cells were increased in cGVHD autoimmune mice

Consistent with a previous study [20], we found that B6 mice transferred with splenocytes from Bm12 mice after 14 days developed a syndrome which closely resembles systemic lupus erythematosus. As illustrated in Figure 1A and B, both mass and cell number of spleens in the cGVHD group was elevated along with higher levels of anti-chromatin IgG and IgG2a antibodies in sera compared with B6 to B6 mice at 14 days after cGVHD induction. It is reported that CD11c^+^ B cell population accumulated in both old female mice and human with autoimmune disease, and might play a direct role in the development of autoimmunity [21]. Our data showed that CD11c^+^ B cells were dramatically increased in the development of cGVHD (Figure 1C).

**Figure 1.**
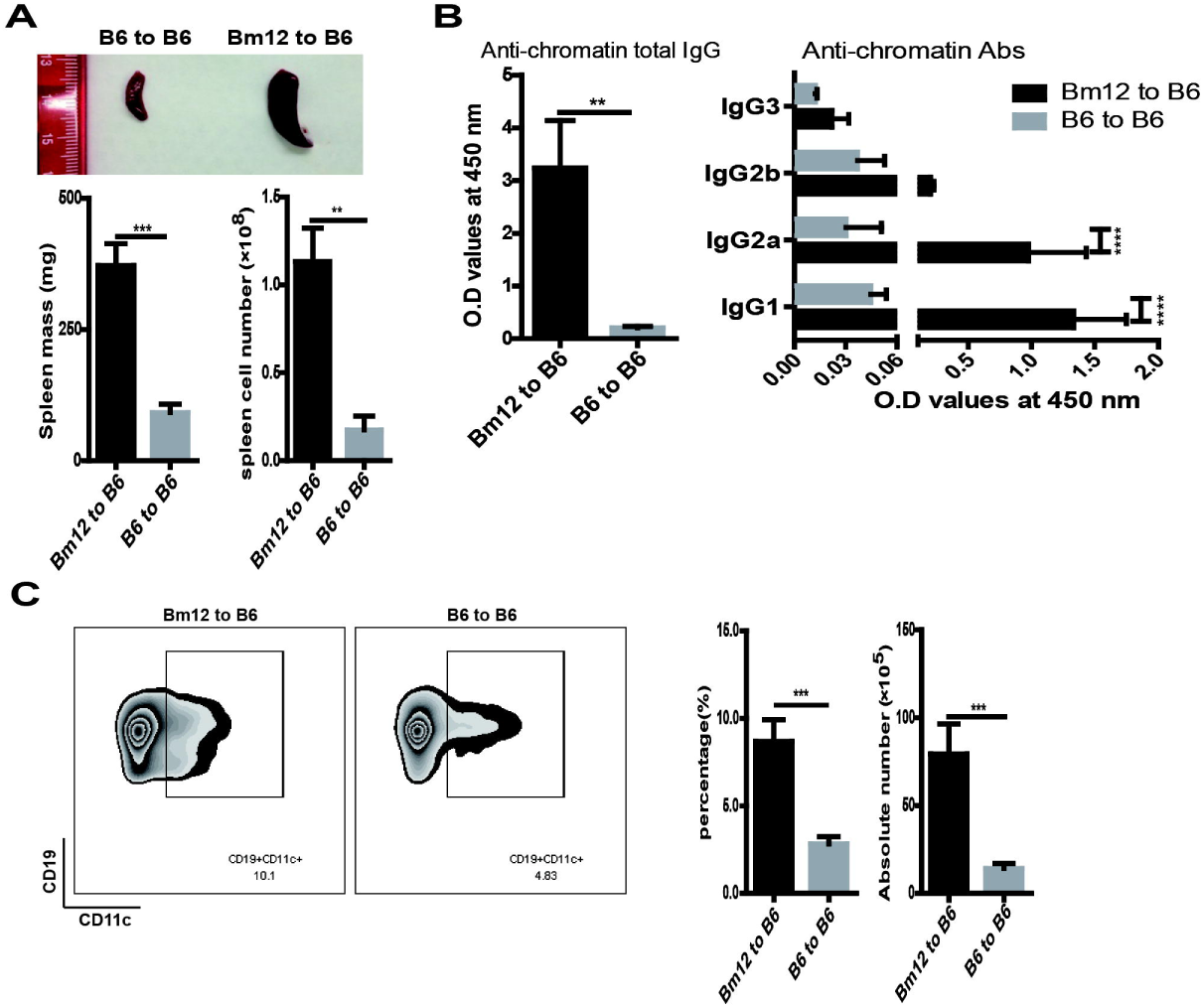
CD11c^+^ B cell was increased in cGVHD. B6 mice (n=5) received an intraperitoneal injection of 5 × 10^^7^ splenocytes from Bm12 or B6, spleens and serum were collected at day 14 for flow analysis and antibodies examination. (A) Weighing and cell counts of spleen were taken 14 days later. The figure shows spleen photos, spleen weight, and absolute number of spleen cells. (B) Serum anti-chromatin total IgG and subtype IgG titers were examined by ELISA. (C) Flow cytometric analysis of CD11c^+^ CD19^+^ cells in total spleen (left) and average percent and absolute number of CD11c^+^ CD19^+^ cells in spleen (right). Values are the mean ± SD. ** = P < 0.01, *** = P < 0.001, **** = P < 0.0001.

Furthermore, CD11c^+^ CD138^+^ plasma cells were also markedly elevated in cGVHD mice (Supplementary Figure 1A and B). Base on above evidence, we speculated that CD11c^+^ B cells were probably involved in the development of lupus by producing autoantibody.

### CD11c^+^ plasma cells produced large amounts of anti-chromatin IgG in vitro

To further investigate the role of CD11c^+^ B cells in cGVHD-induced lupus, we sorted CD11c^+^ CD138^+^, and CD11c^-^ CD138^+^ cells from mice that received Bm12 splenocytes and performed the in vitro functional assay (Fig.2A). As indicated in Figure 2B, CD11c^+^ CD138^+^ cells produced more anti-chromatin IgG antibodies than CD11c^-^ CD138^+^ cells did upon LPS or R848 stimulation, although no statistical significance was observed in the R848 groups.

**Figure 2.**
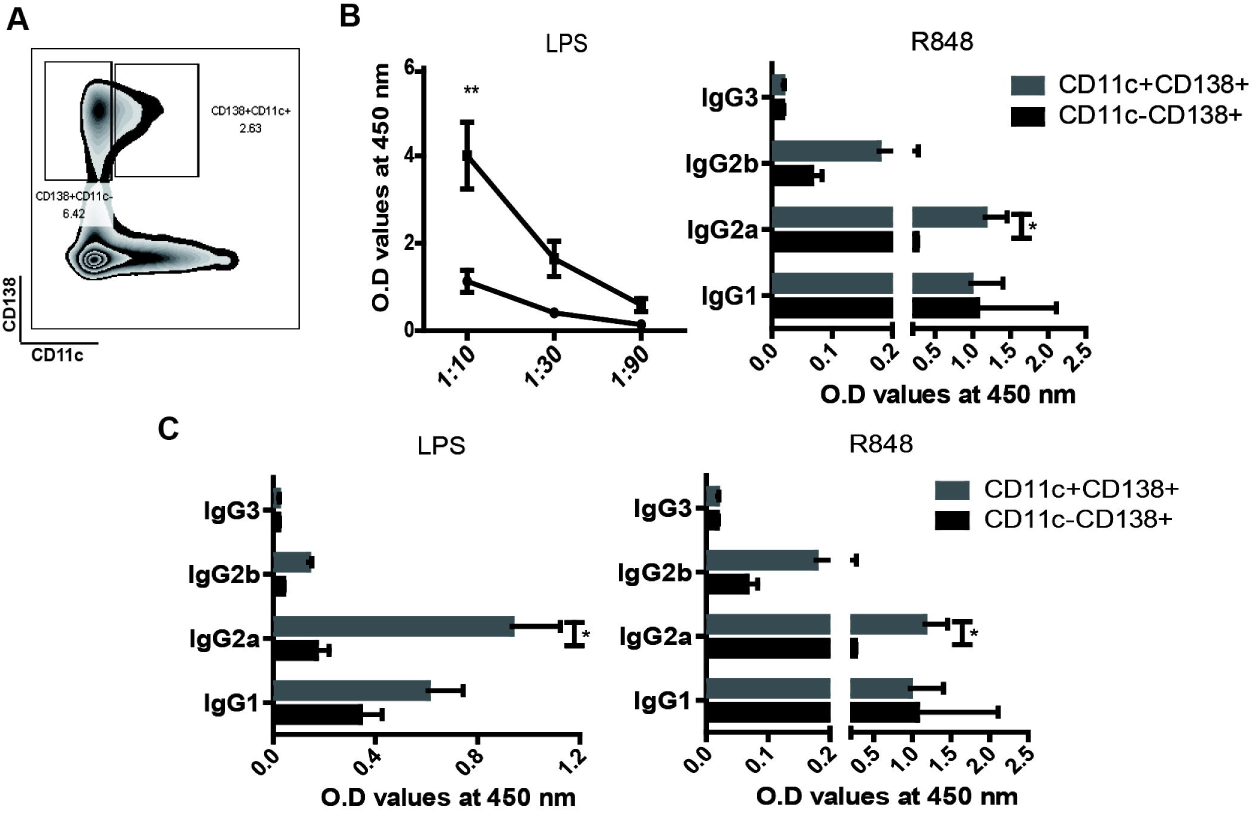
CD11c^+^ plasma cells in cGVHD mice produce anti-chromatin antibodies after stimulation in vitro. (A) CD11c^+^ CD138^+^, and CD11c^−^CD138^+^ cells were sorted from cGVHD mice and cultured for 7 days in the presence of TLR4 (LPS) or TLR7 agonist (R848). Sera anti-chromatin total IgG (B) and anti-chromatin IgG subclass (C) were subsequently measured supernatant by ELISA. (B-C) Bars represent mean (± SD) of 3 independent experiments. * = P < 0.05, ** = P < 0.01.

The effector mechanisms of subclasses of antibodies were distinct due to different constant regions. IgG2a is reported to have the most protective and pathogenic properties among mouse IgG subclasses [27, 28]. Notably, we found that anti-chromatin IgG2a was exclusively produced by CD11c^+^ CD138^+^ plasma cells (Figure 2C).

### Depletion of CD11c^+^ B cells ameliorated anti-chromatin IgG production in vivo

Next, we wanted to know whether depletion of CD11c^+^ B cells in cGVHD mice could reduce the level of anti-chromatin IgG in vivo. To this end, CD11c-DTR mice were transferred with 5 × 10^^^7 splenocytes of Bm12. Then i.p. injected with 100 ng Diphtheria toxin in 200 uL PBS at day 7, day 9 and day 11-post cGVHD induction (Figure 3A). As expected, CD11c^+^ B cells successfully reduced in CD11c-DTR mice induced with cGVHD (Figure3B). Moreover, transient depletion of CD11c^+^ B cells significantly decreased the level of anti-chromatin IgG and IgG2a antibodies in sera of cGVHD mice(Figure 3C), concurrently relieving the splenomegaly. In general, these results demonstrated that CD11c^+^ B cells were critical for anti-chromatin IgG production, both in vitro and in vivo.

**Figure 3.**
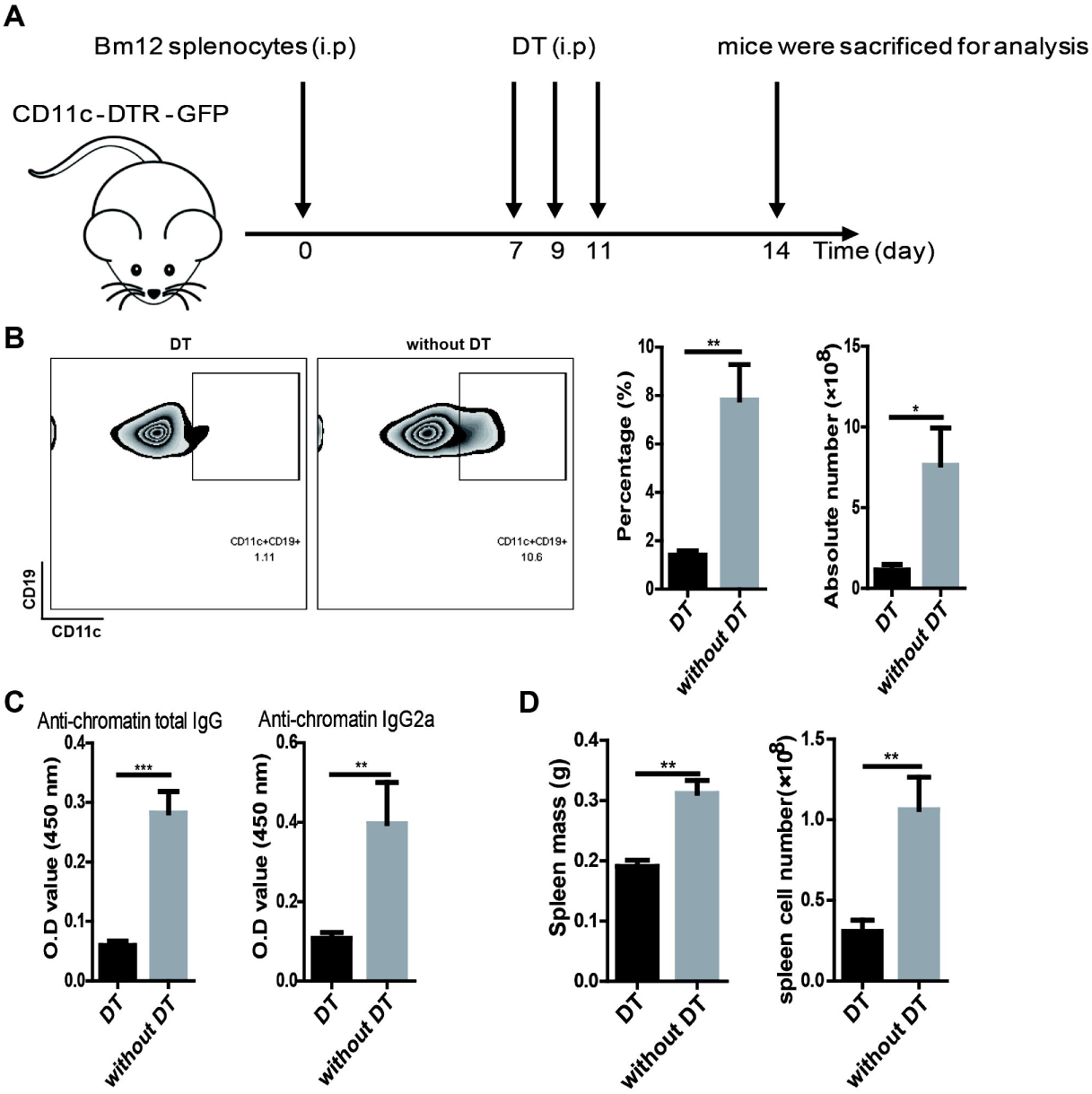
In vivo depletion of CD11c^+^ B cell attenuated the production of anti-chromatin IgG after cGVHD induction. (A) Design of transient depletion CD11c^+^ B in cGVHD study. Mice received 3 every other day intraperitoneal injections of 100ng Diphtheria toxin (DT) in 200 uL PBS (Sigma) (n = 5) or negative control (PBS) (n = 5) for 7 days before the intraperitoneal injection of splenocytes from Bm12. 14 days later, the recipient mice were sacrificed for analysis. (B) Flow cytometric analysis of CD19^+^ CD11c^+^ cells in total spleen with cGVHD induction at 14 days later. (C) Anti-chromatin IgG and IgG2a autoantibody were measured by ELISA in serum of CD11c-DTR mice with cGVHD induction. (D) Spleen weight, and absolute number of spleen cells. Results are representative of 3 independent experiments. In B and C, values are the mean ± SD. * = P < 0.05, ** = P < 0.01, *** = P < 0.001.

### T-bet^+^ CD11c^+^ B cells were significantly increased after cGVHD induction

IgG2a is known as the greatest pathogenic antibody in autoimmune disease [29, 30]. Many studies have demonstrated that T-bet is critical for IgG2a class switching [31, 32] and IgG2a memory response [33]. Recently, it is reported that T-bet drives CD11c^+^ B cells activation to secrete virus-specific IgG2a antibody upon virus infection [34]. Thus, we next investigated the role of T-bet in CD11c^+^ B cell differentiation and anti-chromatin IgG2a production in cGVHD-induced lupus. As shown by qPCR, T-bet expression was remarkably upregulated in spleen B cell of mice obtained splenocytes from Bm12at 14 days. However, no significant change in non B cell between Bm12 to B6 and B6 to B6 (Figure 4A). Furthermore, the mice that received Bm12 splenocytes showed more T-bet^+^ CD11c^+^ CD19^+^ B cells than those received B6 splenocytes (Figure 4B). Meanwhile, CD138^+^ plasma cells were dramatically expanded in cGVHD mice. And surprisingly, above 70% of CD138^+^ cells were IgG2a positive (Supplementary Figure 1B). Interestingly, the fraction of T-bet^+^ CD11c^+^ was significantly increased in CD138^+^ plasma cells (Supplementary Figure 1C). Therefore, these results suggested that T-bet expression may contribute to the production of anti-chromatin IgG2a antibodies in the development of cGVHD.

**Figure 4.**
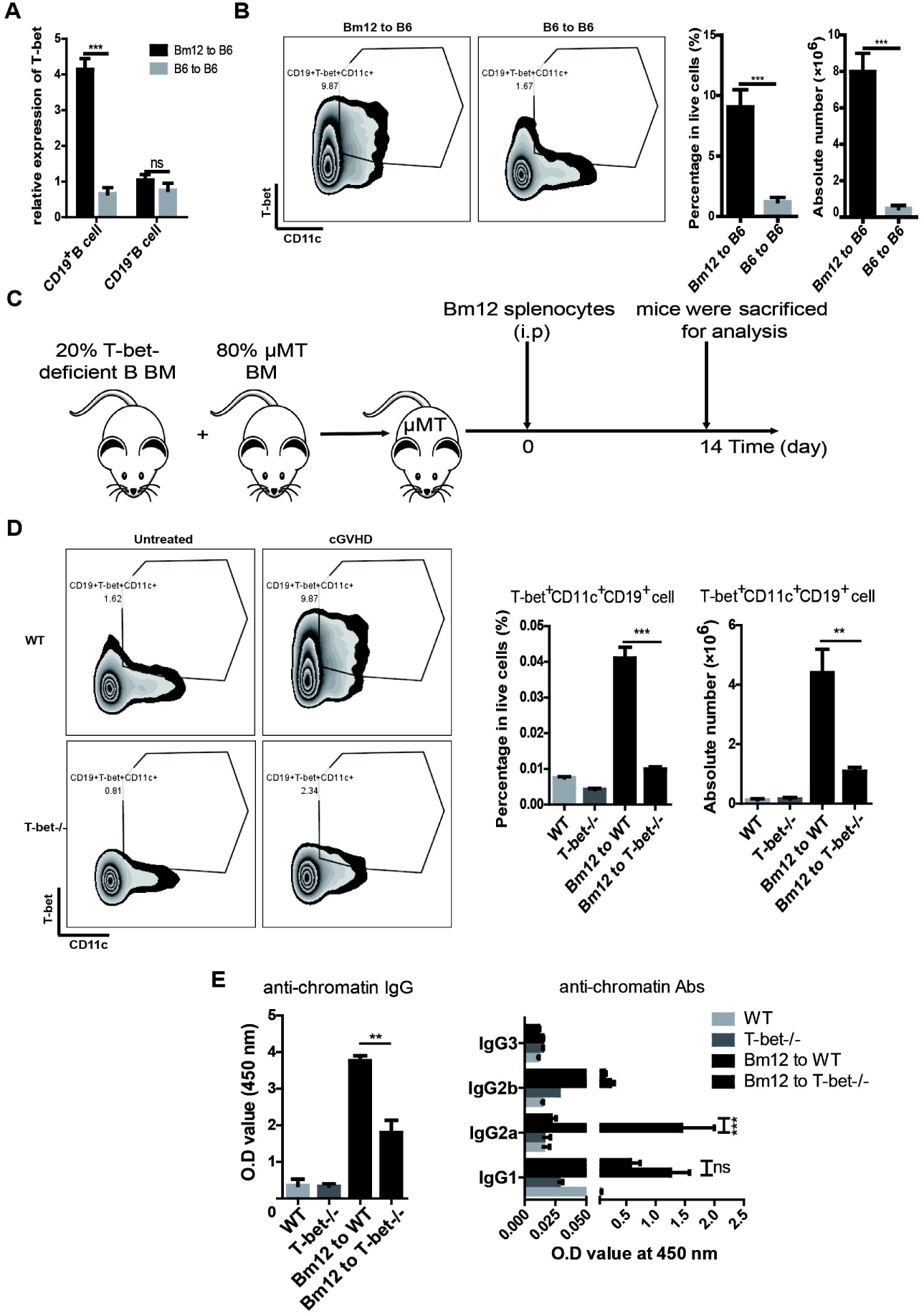
Anti-chromatin IgG production during cGVHD requires T-bet^+^ CD11c^+^ CD19^+^ B cells. (A) CD19^+^ cell and CD19^-^ cell of spleen were isolated from B6 mice transferred with splenocytes of Bm12 or B6 at 14 day for examined T-bet expression. (B) Flow cytometric analysis of T-bet^+^ CD11c^+^ CD19^+^ B cells in total spleen at day 14 with cGVHD induction. (C) Design of T-bet deficient B cell study. B cells of knockout T-bet were transferred into μMT mice (n>5). Then the mice were i.p. injected 5 × 10 Bm12 splenocytes. Spleen and serum were collected at 14 days after cGVHD induction. (D) Flow cytometric analysis of T-bet^+^ CD11c^+^ CD19^+^ B cells in total spleen. (E) ELISA was used for testing the production of sera anti-chromatin IgG. Results in A are representative of 3 independent experiments. Values are the mean ± SD. ** = P < 0.01, *** = P < 0.001, ns = not significant.

### Depletion of T-bet^+^ B cells ameliorated anti-chromatin IgG2a production in vivo

Next, we wondered whether depletion of T-bet^+^ CD11c^+^ CD19^+^ B cells in cGVHD-induced mice could reduce the level of anti-chromatin IgG in vivo. We generated B cell specific T-bet-/- mice by transplanting 20% T-bet-deficient B bone marrow with 80% μMT bone marrow into irradiated μMT mice and then induced the cGVHD model (Figure 4C). As expected, depletion of T-bet^+^ B cells inhibited the expression of CD11c in B cell after cGVHD induction (Figure 4D). Moreover, the level of anti-chromatin IgG and IgG2a was significantly decreased in the absence of T-bet^+^ B cells, yet no changes of anti-chromatin IgG1 production (Figure 4E). Taken together, our data demonstrated that T-bet is critical for CD11c^+^ B cells differentiation and anti-chromatin IgG2a production in cGVHD-induced lupus.

### T-bet^+^ CD11c^+^ CD19^+^ B cells were significantly increased in lupus patients

Next, we investigated the abnormality of T-bet^+^ CD11c^+^ CD19^+^ B cells in lupus patients. As shown in Figure 5A, the frequency of T-bet^+^ CD11c^+^ CD19^+^ B cells was significantly increased in PBMC of lupus patients.

**Figure 5.**
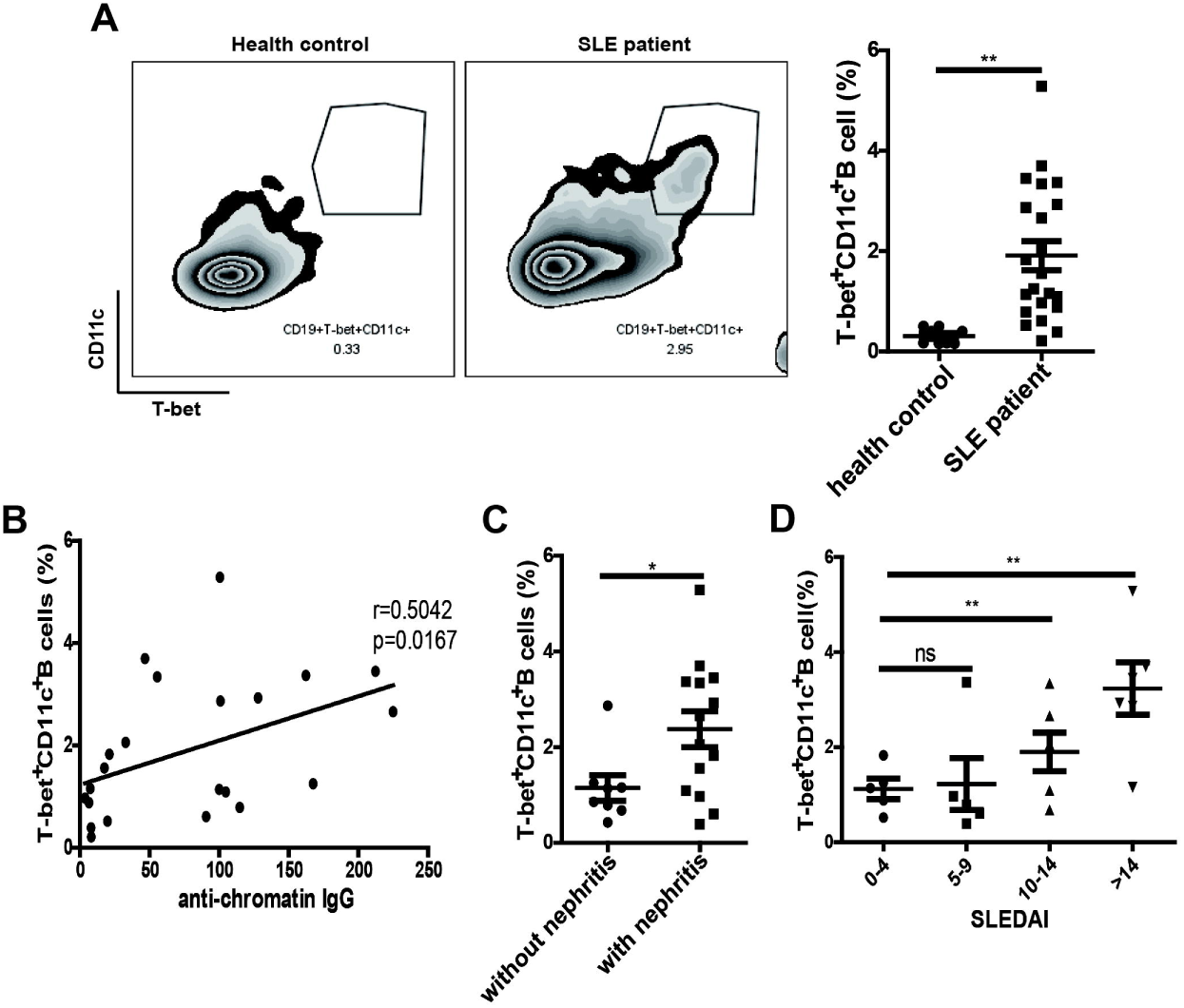
The percentage of T-bet^+^ CD11c^+^ CD19^+^ B cells was elevated and associated with anti-chromatin antibodies in SLE patients. (A) Flow cytometric analysis of T-bet^+^ CD11c^+^ CD19^+^ B cells in peripheral blood from SLE patients (n = 18) and health donors (n = 10). (B) Correlation between the percentage of T-bet^+^ CD11c^+^ CD19^+^ B cells and the level of anti-chromatin antibodies. (C) The percentage of T-bet^+^ CD11c^+^ CD19^+^ B cells exhibited a increasing trend in patients with lupus nephritis (LN; n = 14) relative to patients with no history of LN (n = 8). (D) Patients were classified using the SLE Disease Activity Index (SLEDAI), and the percentage of T-bet^+^ CD11c^+^ CD19^+^ B cells in SLE patients with different disease severity is presented. In A, C, and D, values are the mean ± SD. * = P < 0.05; ** = P < 0.01; in A-D, symbols represent individual subjects.

We then analyzed the relationship between the frequency of T-bet^+^ CD11c^+^ CD19^+^ B cells and autoantibody in the same group of SLE patients. No correlation was observed between the percentage of T-bet^+^ CD11c^+^ CD19^+^ B cells and the presence of anti-dsDNA antibodies and anti-ANA antibodies (Supplementary Figure 3A and B). However, the proportion of T-bet^+^ CD11c^+^ CD19^+^ B cells positively correlated with the serum titer of anti-chromatin IgG antibodies in SLE patients (P = 0.0226, Spearman’s r = 0.5335; Figure 5B). Moreover, we found that the percentage of T-bet^+^ CD11c^+^ CD19^+^ B cells was relatively higher in patients with nephritis than in those without nephritis (Figure 5C). Then we also analyzed the relationship between the percentage of T-bet^+^ CD11c^+^ CD19^+^ B cells and disease activity in the same group of SLE patients. Percentage of T-bet^+^ CD11c^+^ CD19^+^ B cells was higher in patients with moderate or severe disease than in those with inactive disease (Figure 5D). Our data indicated that T-bet^+^ CD11c^+^ CD19^+^ B cells might be responsible for the abnormal levels of anti-chromatin in SLE patients, which might be helpful in the diagnosis and treatment of SLE.

## Discussion

In this manuscript, we characterized an unexpected- CD11c^+^ B cell population appears at high frequency in spleens of lupus-like model (cGVHD), which contributes to the production of anti-chromatin IgG and subclass IgG. Depletion CD11c^+^ B cells attenuated anti-chromatin IgG and subclass IgG2a production, alleviated the manifestation of splenomegaly. Moreover, T-bet is required for CD11c^+^ B cell differentiation. Deletion of T-bet reduced the number of CD11c^+^ B cells and anti-chromatin IgG and IgG2a production in cGVHD-induced lupus. Together, we demonstrated that the expression of T-bet and CD11c in B cells is required for the development of lupus.

Early studies show that T-bet expression in the B lineage promotes class switching to IgG2a [32] and is indispensable for IgG2a memory formation [35]. Recently, Rubtsova et al revealed that T-bet^+^ CD11c^+^ B cells appear at the peak of antiviral response secrete antiviral IgG2a and essential for effective viral clearance [34]. IgG2a is the most potent isotype in mediating the development of lupus [36–38]. In our study, we found that CD138^+^ plasma cells were markedly increased at 14 day in cGVHD mice. And interestingly, above 70% of CD138^+^ cells were IgG2a positive. Moreover, T-bet^+^ CD11c^+^ CD138^+^ cells were significantly increased. The evidence suggests that the expression of T-bet and CD11c might be induced in a small population of B cell in the development of lupus, which might be involved in IgG2a^+^ plasma cell differentiation. However, the molecular regulatory mechanism of T-bet needs to be further explored.

IFNγ was reported to play a pathogenic role in lupus nephritis. Depletion of IFNγ significantly improved renal disease induced by Pristane, with reduction of the production of anti-DNA/chromatin autoantibody [39–41]. Treatment of MRL-Lpr mice with IFNγR/Fc significantly reduced serum levels of IFNγ and autoantibody, consequently improving renal pathology. IFNγ is well known to be able to induce B cell IgG2a class-switching [42]. In vitro study showed that IFNγ and anti-IgM synergistically induced T-bet expression in B cell in a STAT-1-dependent manner to promote IgG2a production [43]. However, it is still unclear how IFNγ triggers B cells to secrete autoantibody in vivo. We noted that the expression of IFNγ was dramatically elevated both in total spleen and in CD4^+^ T cells at day 14-post cGVHD induction (Supplementary Figure 2A). And IFNγ^+^ CD4^+^ T cells were significantly increased in spleens from mice that received Bm12 splenocytes, compared with the control group (Supplementary Figure 2B). Notably, the percentage and absolute number of T-bet^+^ CD11c^+^ CD19^+^ B cells were decreased in IFNGR1-/- mice induced by cGVHD (Supplementary Figure 2C). Thus, we think that IFNγ might induce the expression of T-bet in B cells leading to their differentiation into CD11c^+^ B cells in cGVHD. Further studies are needed to confirm this hypothesis.

Recently, Rubtsov et al reported that CD11c^+^ B cells were expanded only in a handful of SLE patients [21]. Others studies have showed that a resembling population of B cells increased in the peripheral blood of RA patients and some autoimmune individuals with common variable immunodeficiency [44–46]. In this study, we demonstrated that T-bet^+^ CD11c^+^ CD19^+^ B cell population was significantly increased in the peripheral blood of SLE patients compared with healthy control. Interestingly, the percentage of T-bet^+^ CD11c^+^ CD19^+^ B cells in blood was associated with the sera level of anti-chromatin, and elevated in SLE patients with lupus nephritis and moderate or severe disease, with no collection with anti-ANA antibodies or anti-dsDNA antibodies, implying the specificity for anti-chromatin. Furthermore, we also detected that T-bet^+^ CD11c^+^ CD19^+^ B cells were presented at higher proportion in patients with severe or moderate SLE than in those with inactive SLE. These data suggest that T-bet^+^ CD11c^+^ CD19^+^ B cells might serve as a biomarker for clinical diagnosis and treatment for lupus patients.

In summary, we have demonstrated that a newly discovered population of B cell, T-bet^+^ CD11c^+^ CD19^+^ B cell is associated with the titer of anti-chromatin in lupus patients and directly involved in secretion of autoantibody in cGVHD mice. Targeted depletion of CD11c or T-bet efficiently reduced the production anti-chromatin. These findings highlight that T-bet^+^ CD11c^+^ CD19^+^ B cells might be a potential therapeutic target of lupus.

